# A robust measure of metacognitive sensitivity and confidence criteria

**DOI:** 10.1101/2023.09.12.557480

**Authors:** Derek H. Arnold, Mitchell Clendinen, Alan Johnston, Alan L.F. Lee, Kielan Yarrow

**Affiliations:** School of Psychology, The University of Queensland; School of Psychology, The University of Nottingham; Department of Psychology, Lingnan University, Hong Kong; School of Psychology, City University London

**Keywords:** Perceptual metacognition, Confidence, Signal Detection Theory

## Abstract

Humans experience feelings of confidence in their decisions. In perception, these feelings are typically accurate – we tend to feel more confident about correct decisions. The degree of insight people have into the accuracy of their decisions is known as metacognitive sensitivity. Currently popular methods of estimating metacognitive sensitivity are subject to interpretive ambiguities because they assume that humans experience normally-shaped distributions of different experiences when they are repeatedly exposed to a single input. If, however, people have skewed distributions of experiences, or distributions with excess kurtosis (i.e. a distribution containing greater numbers of extreme experiences than is predicted by a normal distribution), calculations can erroneously underestimate metacognitive sensitivity. Here, we describe a means of estimating metacognitive sensitivity that is more robust against violations of the normality assumption. This improved method relies on estimating the precision with which people transition between making categorical decisions with relatively low to high confidence, and on comparing this with the precision with which they transition between making different types of perceptual category decision. The new method can easily be added to standard behavioral experiments. We provide free Matlab code to help researchers implement these analyses and procedures in their own experiments.

**Public Significance Statement:** Signal-detection theory is one of the most popular frameworks for analysing data from experiments of human behaviour – including investigations of confidence. The authors demonstrate that if a key assumption of this framework is inadvertently violated, analyses of confidence can lead to unwarranted conclusions. They develop a new and more robust measure of confidence.

## Introduction

People experience levels of confidence when making decisions (Fleming et al., 2012; Yeung & Summerfield, 2012). In studies of perception, we can verify that these feelings are typically accurate, with higher levels of confidence experienced for better decisions (De Martino et al., 2013; Keane et al., 2015; Li et al., 2014; Peters et al., 2017). This is indicative of a level of insight into the quality of our decisions – known as metacognitive sensitivity (Maniscalco & Lau, 2012, 2016). When confidence appears to have been informed by the same information as our decisions, people are said to be metacognitively ideal (Fleming and Lau, 2014; Maniscalco & Lau, 2012, 2016). However, when confidence appears to have been shaped by different information, people are understood to have inferior insight into their decisions and are said to be metacognitively insensitive (Fleming and Lau, 2014; Maniscalco & Lau, 2012, 2016).

A range of methods have been used to estimate metacognitive sensitivity (or rather the degree of metacognitive insensitivity). A previously popular approach was to assess correlations between the accuracy of perceptual decisions and confidence ratings (e.g. Nelson, 1984). This approach, however, had a serious shortcoming. Such measures can confound confidence bias – an individuals’ propensity to report that they are confident, with the desired measure of metacognitive sensitivity. Two people can have equal levels of metacognitive sensitivity, have equally heightened feelings of confidence when they are making a well-informed relative to an ill-informed decision, but have different levels of willingness to report being confident. This confidence bias can obscure an individuals’ true level of metacognitive sensitivity (Fleming and Lau, 2014; Maniscalco & Lau, 2012, 2016).

More recent attempts to measure metacognitive sensitivity have relied on an extrapolation of the Signal-Detection-Theory (SDT) framework (Fleming & Lau, 2014; Maniscalco & Lau, 2012). In addition to the standard SDT framework, these approaches assume people use criteria to demarcate when a decision is regarded as having evoked a relatively low or a relatively high feeling of confidence (see Figure 1). With this added assumption, two sets of estimates of perceptual bias and sensitivity can be calculated from a common dataset. Traditional estimates of sensitivity (d’) and bias can be calculated from perceptual category decisions (Green & Swets, 1966). Researchers can also backwards infer a second estimate of sensitivity and bias, from analyses that are informed by the proportions of correct (e.g. Figure 1, bottom panel, light red and blue shaded regions) and incorrect (e.g. Figure 1, bottom panel, dark purple shaded regions) high-confidence decisions (Fleming & Lau, 2014; Maniscalco & Lau, 2012). The key statistic this process delivers is meta d’, which is regarded as an estimate of metacognitive sensitivity when it is expressed relative to d’ estimates, by either calculating difference scores (meta d’ – d’) or ratios (meta d’ : d’). The standard finding is that meta d’ estimates fall short of d’ estimates (e.g. Maniscalco et al., 2016; Rausch et al., 2015), which is regarded as evidence that confidence ratings have been shaped by different information relative to perceptual decisions.

**Figure 1.**
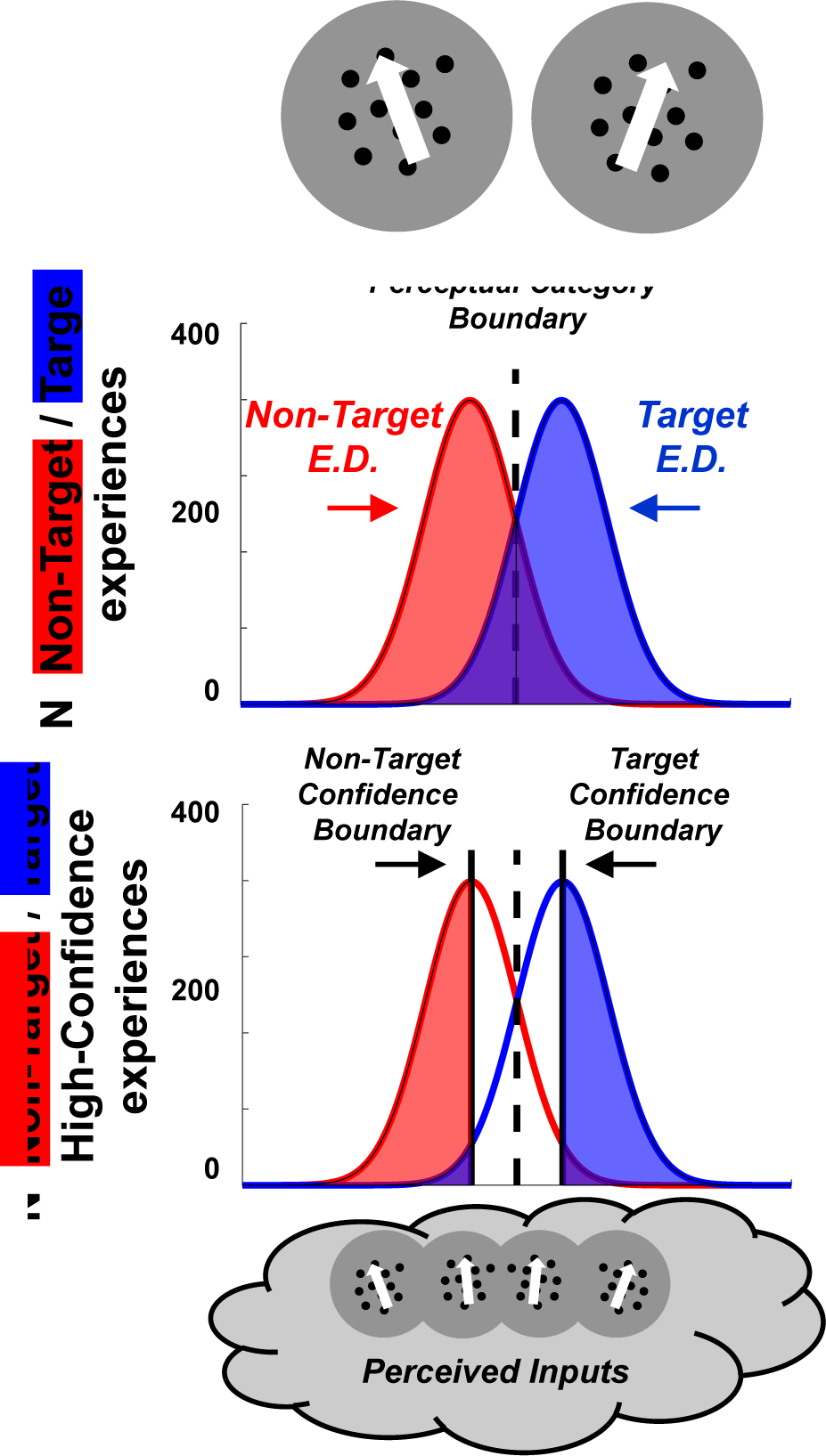
Graphic depicting the assumptions underlying popular extensions of the SDT framework, used to analyze confidence. **Above**, a standard SDT decision space is depicted, with a non-target (red) and a target (blue) experiential distribution depicted, and a central perceptual category boundary (bold black dotted vertical line). **Below**, a non-target-(left bold vertical line) and a target confidence criterion (right bold black vertical line) are assumed to demarcate experiences that are classified with low or with high confidence. These criteria are applied to experiences elicited by both non-target and by target presentations. Using this logic, researchers can backwards infer how separated target and non-target experiential distributions must be, to be consistent with the proportions of *correct* (light red and light blue shaded regions) and *incorrect* (dark purple shaded regions) high-confidence categorizations. This estimate is known as meta-d’.

SDT-based analyses of confidence are regarded as an improvement on correlations of confidence with decisional accuracy (e.g. Nelson, 1984), as they provide independent estimates of metacognitive sensitivity and confidence bias (Fleming & Lau, 2014; Maniscalco & Lau, 2012). They can, however, encourage erroneous conclusions if they are inadvertently informed by experiential distributions (E.D.s) that are non-normally shaped – either because the E.D. is skewed (see Arnold et al., 2023), or because the E.D. has fatter tails (a greater number of extreme experiences, called excess kurtosis) than is predicted by a normally-shaped E.D. (see Arnold et al., 2023; Miyoshi et al., 2022). This is a serious consideration for researchers, as there is good evidence that experiential distributions in perception are subject to both excess kurtosis (e.g. Acerbi et al., 2012; Anderson, 2014; Bays, 2016; Jabar & Anderson, 2015) and to localized skews (e.g. Appelle, 1972; Girshick et al., 2011; Storrs & Arnold., 2015b). Moreover, it has been established that current methods of assessing if E.D.s are likely to have been normally shaped can be insensitive to deviations from normality that are sufficient to cause meta d’ to be underestimated (see Arnold et al., 2023). So, there is an interpretive ambiguity. Any meta d’ - d’ difference could either be due to confidence having been shaped by different information relative to perceptual decisions, or it could have arisen because decisions were informed by a common set of non-normally shaped E.D.s.

Given these considerations, we were motivated to develop a more robust means of estimating metacognitive sensitivity. We have found this can be achieved in experiments where people decide if inputs belong to one of two categories (a binary forced choice task, a standard procedure in studies of perception). We estimate the precision with which participants transition from making low-to high-confidence perceptual category decisions, and compare these estimates to the precision with which they transition from predominantly making one type of perceptual category decision to another (see Figure 2). In both cases cumulative Gaussian functions are fit to data, and slopes at the inflection points of fitted functions are taken as estimates of the precision of the different types of judgment (about perception and confidence). Additionally, the separation of distributions describing transitions in confidence from a central distribution that describes transitions between perceptual categories, provides an estimate of a participants’ criterion for reporting that they have a relatively high-level of confidence (see Figure 2).

**Figure 2.**
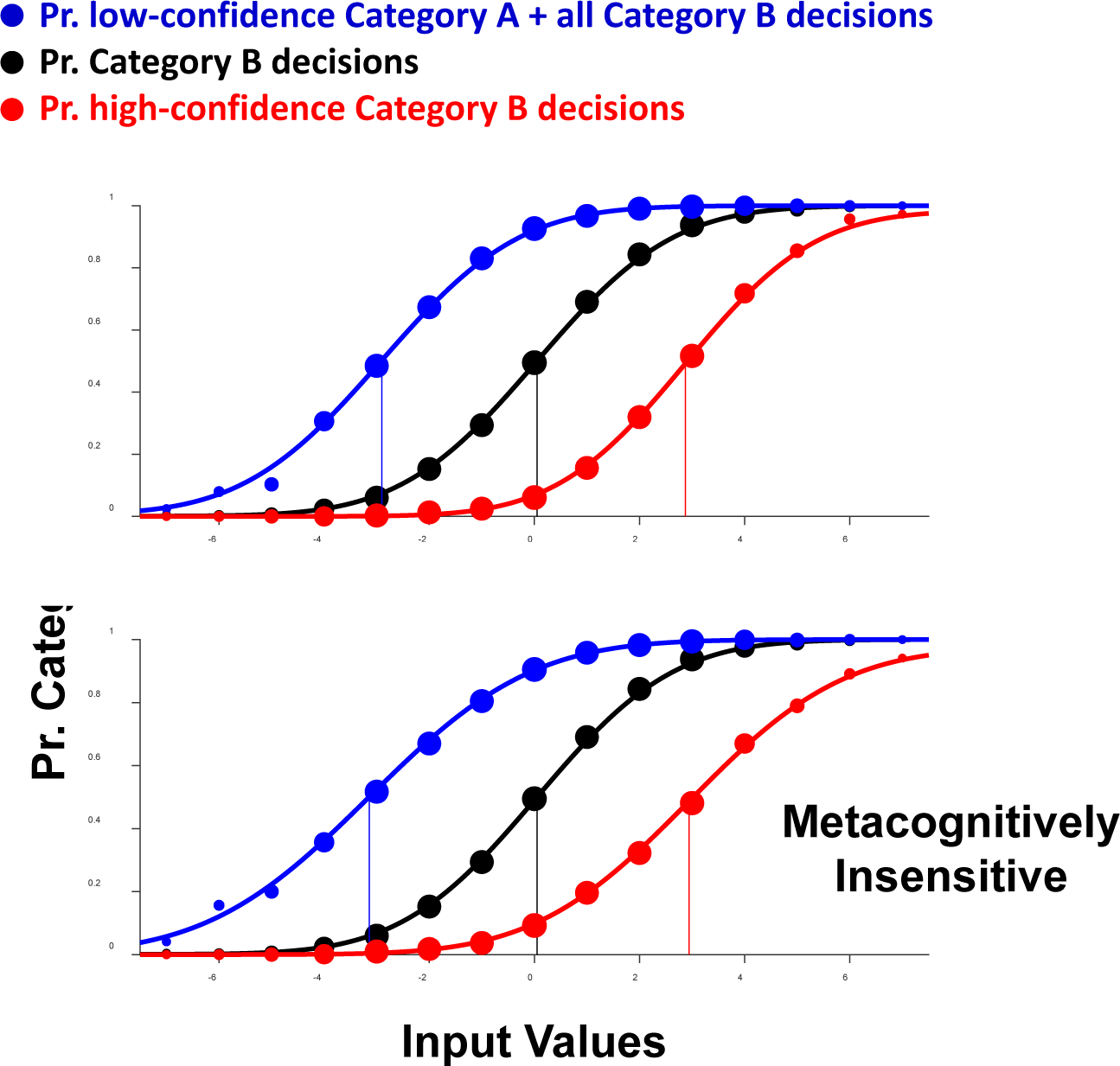
Cumulative gaussian functions fit to **1)** the proportion of simulated trials resulting in either low-confidence category A perceptual decisions or in any category B perceptual decision (**blue data**), **2)** the proportion of simulated trials resulting in any category B perceptual decisions (**black data**), or **3)** the proportion of high-confidence category B decisions (**red data**). The average slope at the inflection points of the **blue** and **red** functions are regarded as estimates of the precision of confidence ratings, whereas the slope at the inflection point of the **black** functions are regarded as an estimate of the precision of perceptual judgments. A metacognitively ideal (top) and metacognitively insensitive (bottom) observer are depicted.

To clarify, imagine participants are completing an experiment where they must decide if oriented lines are tilted left or right of vertical, and they must also express some level of confidence in these decisions. In a standard experimental design, we could repeatedly present inputs that have a range of different physical tilts, and plot the proportion of times the participant reported that each had been tilted right of vertical (see black data, Figure 2a). We can then fit a cumulative Gaussian function, and regard the X-axis value of the inflection point of that function as an estimate of the subjective perceptual category boundary (i.e. the physical tilt at which the participant transitioned from predominantly reporting that inputs were left tilted to predominantly reporting that they were right tilted). This, of course, is subject to bias, which can be a desirable feature as we often want to manipulate and measure bias in perception research (e.g. Clifford et al., 2000; Gibson & Radner, 1937; Regan & Beverly, 1985; Webster, 2015). The slope of the fitted function at the inflection point can be regarded as an estimate of the precision of the perceptual judgments. The steeper the function, the more precise the perceptual judgments, as when a participant reliably categorizes input tilts across many trials, they will transition from predominantly making one category of perceptual decision (i.e. left tilted) to predominantly making the other (right tilted) across a small range of physical tilts.

Once an experimental session has finished, we can regard trial confidence ratings as having been low or high for a participant by comparing each rating to the average confidence rating for that participant across all trials. This can be done for confidence ratings taken from settings along a continuous scale, or for confidence ratings selected from Likert scale options. We can then plot two additional functions, that each describe transitions in confidence. The first (see blue data, Figure 2) plots the proportion of trials on which the participant has either made a Category A decision with low confidence (i.e. a low-confidence left tilt decision), or any Category B decision (i.e. a right tilt decision, regardless of confidence). The second function (see red data, Figure 2) plots the proportion of trials on which the participant has made a Category B perceptual decision with high confidence. In both cases, a cumulative Gaussian function is fit to data, and the coincidence of its inflection point with the X-axis can be regarded as an estimate of the confidence boundary for that category of perceptual decision. The horizontal separation of these two confidence boundaries from the central perceptual category boundary provides a measure of the participants’ confidence criterion (in this case, the degree of tilt from subjective vertical that the participant required before they would endorse a perceptual category decision with a relatively high level of confidence). The slope of these functions at their inflection points can be regarded as an estimate of the precision of confidence ratings.

If confidence ratings are informed by the same information as perceptual decisions, and the criteria people use to categorize their feelings of confidence (as relatively low or high) are equally stable relative to the perceptual category criterion, then estimates of the precision of confidence ratings and perceptual category decisions should be equal. If, however, confidence ratings are shaped by different information (either from processes that generate our feelings of confidence, or from processes that are used to apply confidence criteria), then estimates of the precision of confidence ratings should fall short of estimates of the precision of perceptual decisions. The degree to which this is true can be regarded as an estimate of metacognitive inefficiency.

To compare our new approach with existing SDT-based analyses of confidence, we first present analyses of data from simulated experimental sessions. In a first set of simulations, we assess how the two methods are impacted by E.D.s that have different levels of excess kurtosis. Then, in a second set of simulated experiments, we examine the impact of skewed E.D.s. In both cases we show that our new method is more robust to deviations from normally shaped E.D.s.

We then report on the results of a series of 3 behavioural experiments. These serve to validate our new method of estimating metacognitive sensitivity in experiments with human participants. The first two of these re-examines the impact of the range of direction signals on confidence when people make judgments about the average direction (Spence et al., 2016). The first study has people report on confidence using a continuous scale (Experiment 1), the second has people report on confidence using a Likert scale (Experiment 2). In both cases we find that estimates of metacognitive sensitivity are constant across experimental conditions (with different ranges of direction signal), but when there is a greater range of direction signals people are more conservative when reporting on confidence (i.e. they require a greater difference in average direction before they will indicate having a relatively high-level of confidence in their direction decisions). In Experiment 3 we examine the impact of having people guess, before stimulus presentations, about what test direction will be presented. We find this both decreases metacognitive sensitivity for judgments regarding directions that are guess consistent, and it causes people to be relatively more conservative when reporting on confidence in decisions that are guess inconsistent.

## Simulated Experiments 1a and 1b: Excess Kurtosis

We tested simulated observers with different levels of kurtosis in their experiential distributions, and who had different high confidence criteria. The E.D.s of these simulated observers were either normally shaped (an excess kurtosis of 0), or they had a greater level of kurtosis (fatter tails) than a normally shaped distribution (excess kurtosis levels of 0.2 and 0.4). In all cases, the S.D. of E.D.s was set to 2, and E.D. values were referenced against a perceptual category criterion of 0 (to simulate the performance of unbiased observers). For each level of kurtosis simulated observers with a range of 51 different confidence criteria (ranging from 2.5 to 6) were simulated. These criteria represent the absolute difference in orientation (+/- from the 0 perceptual category boundary) required to elicit a high confidence (1) response. All lesser differences evoked a low confidence (0) response. So, in each experiment we simulate 153 observers (3x kurtosis levels x 51 high confidence criteria). Matlab code to implement these simulated experiments is provided as Supplemental Code #2.

In Experiment 1a we simulate metacognitively ideal participants, with no additional confidence noise. If analyses accurately assess metacognitive sensitivity (i.e. if they provide a metric that indicates if confidence and perceptual decisions have been informed by the same information) then ratios that describe the precision of confidence judgments to the precision of perceptual judgments should equal 1. In Experiment 1b confidence noise is simulated by randomly varying the confidence criterion +/- from its nominal value on a trial-by-trial basis by up to +/- 75%. For these simulated observers, ratios that describe the precision of confidence judgments to the precision of perceptual judgments should have a value less than 1.

Experiments 1a and 1b simulate a forced-choice perceptual categorization task, where on each trial an E.D. value is elicited by a test input, which is then categorized as having been left (E.D. values <0) or right (E.D. values >=0) tilted. For confidence ratings the sign (+/-) of E.D. values is discarded. Confidence ratings are then either set to 0 (when unsigned E.D. values are < the high confidence criterion) or to 1 (when unsigned E.D. values are >= the high confidence criterion). This simulates an experiment where participants make perceptual category decisions and commit to binary subjective confidence ratings (low vs high confidence). In both experiments experimental sessions consist of 20,000 simulated individual trials.

## Slope tests of metacognitive sensitivity

### Sampling procedure and response categorizations

To implement our slope-based tests of metacognitive sensitivity, an appropriate range of test inputs needs to be sampled. To achieve this, 5 staircase procedures are interleaved (Levitt, 1971). The step size for each procedure here was 1. Two staircase procedures were designed to sample inputs that resulted in ∼90% correct high-confidence Category A and Category B perceptual decisions. This was achieved by stepping input values away from the perceptual category boundary whenever 2 or more of the past 10 staircase responses were either incorrect or low-confidence. Otherwise, test input values were stepped toward the perceptual category boundary. These two staircases procedures were each sampled on 5% of session trials.

A further two staircase procedures were designed to sample confidence boundaries for each perceptual category. Test values for these procedures were stepped toward the perceptual category boundary whenever a response was consistent with the targeted category and the subjective confidence rating was high. Otherwise test values were stepped away from the perceptual category boundary. These staircases were each sampled on 30% of experimental session trials.

The final staircase procedure was designed to sample inputs at the perceptual category boundary. Test values for this procedure were stepped away from the perceptual category of response on each trial. Category A responses resulted in values being stepped toward Category B values, and vice versa. This staircase was also sampled on 30% of experimental session trials.

In our simulated experimental sessions, each potential test input was associated with a pre-calculated array of E.D. values. These were sampled in turn whenever that test input value was sampled by a staircase. Each E.D. had a mean equal to the test input value, a S.D. of 2, and a skew of 0. There were 3 sets of simulated observers. For one the kurtosis of E.D. values was 3 (consistent with a normally-shaped distribution, and therefore with an excess kurtosis of 0). In the other sets excess kurtosis values of 0.2 and 0.4 were sampled.

## Results

Ratio values that describe the precision of confidence judgments divided by the precision of perceptual judgments are plotted in Figure 3a, as a function of confidence criterion values. Note that the precision of confidence judgments is taken as the average of the two fitted confidence functions (see Figure 2). These estimates were calculated from experimental sessions that simulated metacognitively ideal participants, so ratios should approximate a value of 1. This was true for all confidence criterion values, and for each level of excess kurtosis (0 – black datapoints, 0.2 – blue data points, and 0.4 – red data points). Nearly all ratios fell within +/-5% of a ratio of 1 (see the green shaded region), and the few exceptions all fee within +/-10% of a ratio of 1. This shows that our technique for estimating metacognitive sensitivity is robust to analyses that are informed by E.D.s with excess kurtosis.

**Figure 3.**
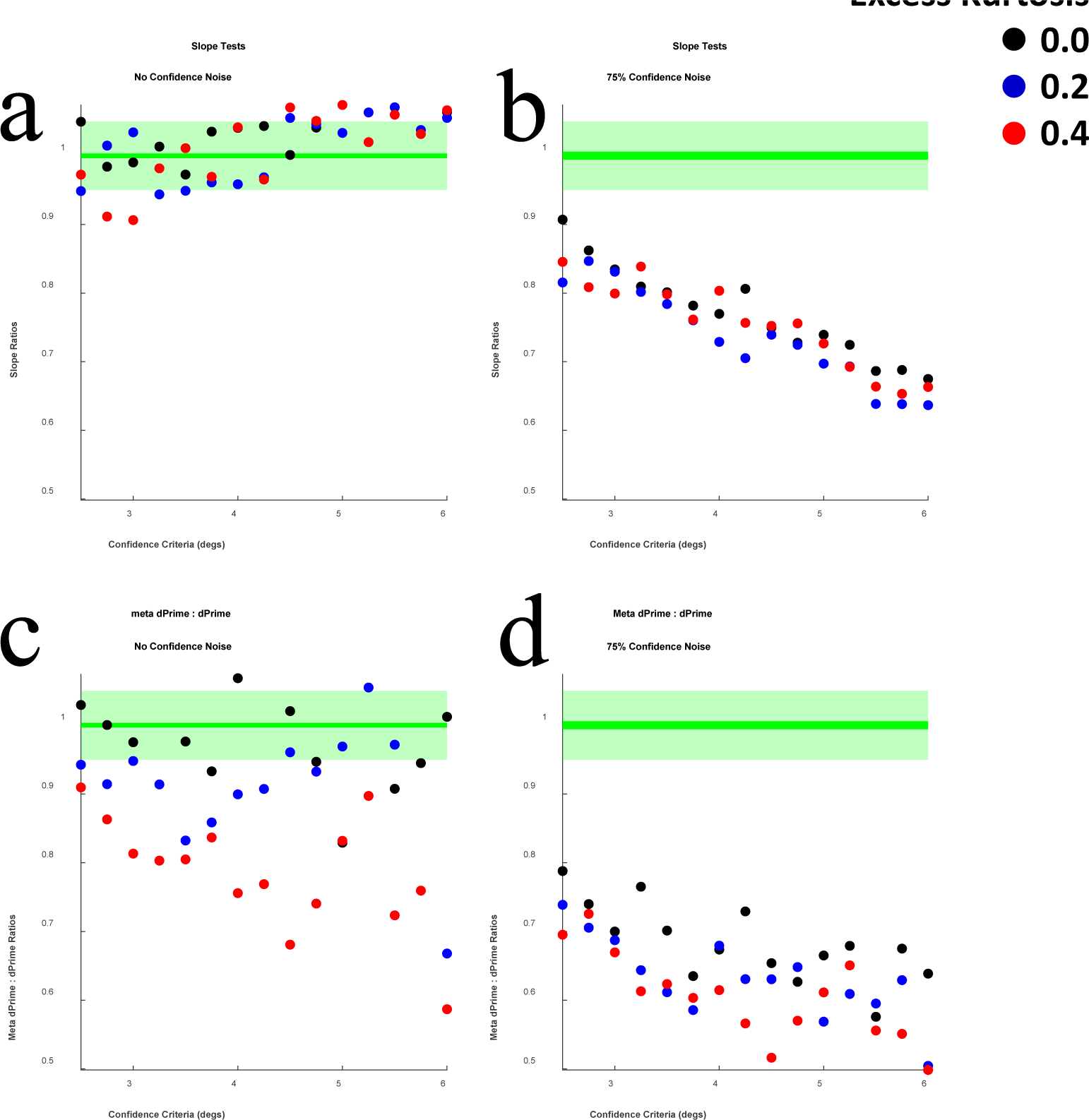
**a)** Scatterplot of confidence : perceptual slope ratios (Y axis) and confidence criterion test values (X axis). These data relate to simulated participants who were metacognitively ideal. The bold horizontal green line depicts ratios of 1, and the green shaded region depicts ratios between 0.95 and 1.05. Black data points depict ratios calculated from sessions that sampled normally-shaped E.D.s, blue data points depict ratios calculated from sessions that sampled E.D.s with an excess kurtosis of 0.2, and red data points depict ratios calculated from sessions that sampled E.D.s with an excess kurtosis of 0.4 **b)** Details are as for Figure 3a, but these data relate to simulated participants who were metacognitively insensitive – due to trial-by-trial confidence criterion noise. **c)** Details are as for Figure 3a, but these data relate to meta d’ : d’ ratios (Y axis) calculated for simulated participants who were metacognitively ideal – in that their perceptual and confidence decisions were informed by a common source of information. **d)** Details are as for Figure 3c, but these data relate to simulated participants who were metacognitively insensitive – due to trial-by-trial confidence criterion noise.

Data plotted in Figure 3b are similar to data plotted in Figure 3a, but these data were calculated from simulated participants who were metacognitively insensitive. This was simulated by randomly varying the nominal confidence criterion value by up to +/- 75% on a trial-by-trial basis. These slope ratios should therefore have a ratio <1. As can be seen in Figure 4b, slope ratios were all less than 1, with all ratios >10% below a ratio of 1. These data show that our test of metacognitive sensitivity can detect cases of genuine metacognitive insensitivity, caused by confidence ratings being shaped by a different source of information relative to perceptual decisions.

**Figure 4.**
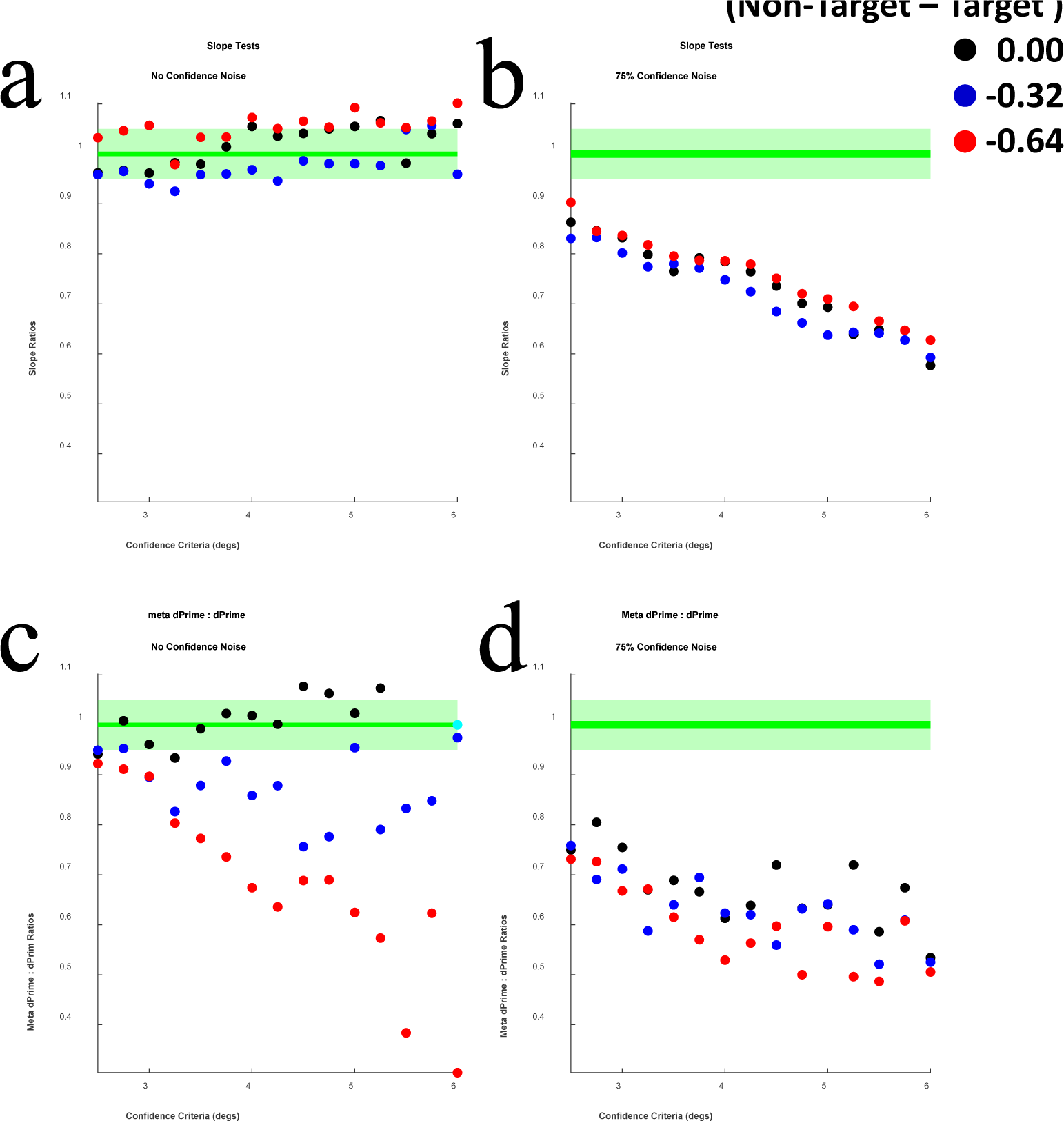
**a)** Details are as for Figure 3a, with the exception that blue data points depict ratios calculated from sessions that sampled E.D.s with a skew difference of 0.32, and red data points depict ratios calculated from sessions that sampled E.D.s with a skew difference of 0.64 (see main text for further explanation). **b)** Details are as for Figure 4a, but these data relate to simulated participants who were metacognitively insensitive – due to trial-by-trial confidence criterion noise. **c)** Details are as for Figure 4a, but these data relate to meta d’ : d’ ratios (Y axis) calculated for simulated participants who were metacognitively ideal. **d)** Details are as for Figure 4c, but these data relate to simulated participants who were metacognitively insensitive – due to trial-by-trial confidence criterion noise.

**Figure 4.**
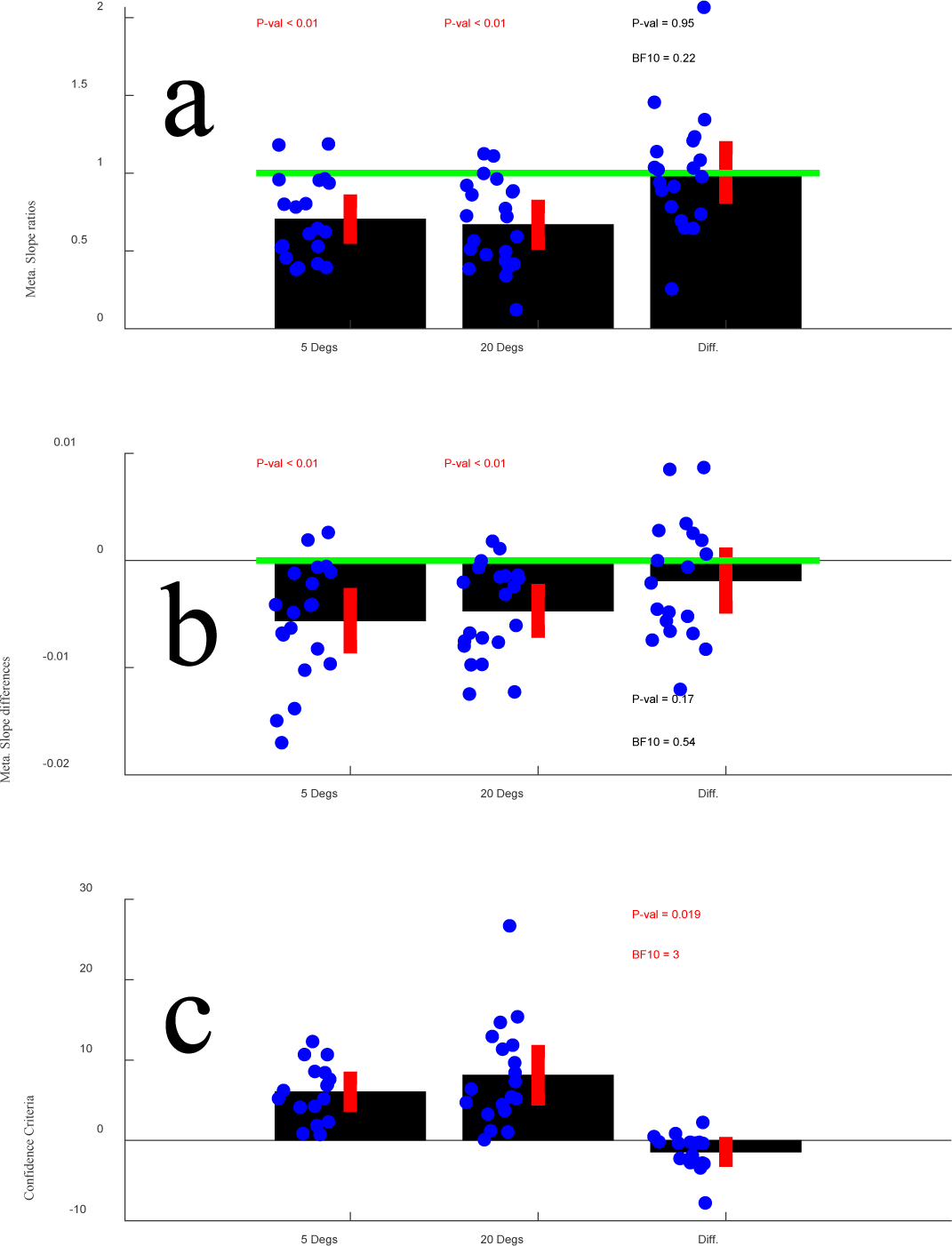
Results of Experiment 3. **a)** Bar plots, depicting Confidence to Perceptual function fit slope ratios, for RDKs with a uniform range of direction signals +/- 5° from the mean (5 degs) and +/- 20° from the mean, along with values of 1 - the difference between these two ratios (Diff). Blue data points depict individual ratios, and red vertical bars depict +/- 2SEM from the group average. The green horizontal line marks a ratio of 1, which is consistent with metacognitively ideal performance (+/- 5 and 20°), and with there being no difference in metacognitive sensitivity across these two conditions (Diff). **b)** Details are as for Figure 4a, but for data relating to slope differences. Here, differences of 0 are consistent with metacognitively ideal performance, and with there being no differences across the two conditions. **c)** Details are as for Figure 4b, but for data relating to confidence criteria. These data do not speak to metacognitive sensitivity.

## Simulated Experiment 1b: SDT-based analyses of confidence and excess kurtosis

We conducted this experiment to compare our new test of metacognitive sensitivity to a currently popular SDT-based analysis (Maniscalco & Lau, 2012). To this end, we simulate a SDT-based experiment, informed by E.D.s that have equal levels of kurtosis relative to the E.D.s that informed Experiment 1a.

## Sampling procedure and response categorizations

Details are as for Experiment 1a, with the following exceptions.

Just 2 test inputs are sampled. An E.D. of 10,000 values is calculated for each of these. As for Experiment 1a, the S.D.s of these is set to 2, and mean values are set to +/- 1. The E.D. values are referenced against a perceptual category criterion of 0 – to simulate an unbiased observer. Also, as per Experiment 1a, a range of 51 different confidence criteria (from 2.5 to 6) are sampled. E.D. values are referenced against these criteria, to determine if the perceptual decision will be endorsed with a low (0) or a high-level (1) of confidence.

For each simulated trial, E.D. values are categorized as having elicited a high-confidence Category A (left tilt) response, a low-confidence Category A response, a low-confidence Category B (right tilt) response, or a high-confidence Category B response – with confidence ratings evaluated against the relevant high confidence criterion. This enables us to implement a popular SDT-based analysis of confidence (Maniscalco & Lau, 2012), which we use to calculate d’ and meta d’ estimates.

## Results

Meta d’ to d’ ratios are plotted in Figure 3c, for experimental sessions that simulate a metacognitively ideal participant. As the participant is metacognitively ideal, ratios should cluster about a value of 1. As can be seen in Figure 3c, this is only true in a broad sense when the E.D.s informing data analyses were normally shaped (black data). When analyses were informed by E.D.s that had excess kurtosis (of 0.2 – blue data, or 0.4 – red data) all ratios were > 5% below a ratio of 1, with a clear increase in the magnitude of reduction for increasingly large confidence criterion values. These data show that a SDT based analysis of confidence can encourage erroneous conclusions when analyses are informed by E.D.s that are non-normally shaped, due to having excess kurtosis relative to a normal distribution. Here the same information has informed d’ and meta d’ calculations, and yet the results would encourage a researcher to conclude that the participant had been metacognitively insensitive – that confidence ratings had been shaped by an additional source of information relative to perception.

Meta d’ to d’ ratios are plotted in Figure 3d, for experimental sessions that simulate a metacognitively insensitive participant. Recall that this was simulated by randomly varying the nominal confidence criterion value by +/- 75% on a trial-by-trial basis. As can be seen in Figure 3d, these slope ratios are all substantially less than 1, and there is a clear decrease in ratio for increasingly large confidence criterion values. These data show that the simulated participants would be identified as metacognitively insensitive by this process. However, these classifications would not be truly diagnostic of metacognitive insensitivity, as the same process has classified many metacognitively ideal simulated participants as metacognitively insensitive when E.D.s had excess kurtosis (see Figure 3c).

## Simulated Experiment 2a: Slope-based tests of metacognitive sensitivity and skew

### Sampling procedure and response categorizations

All details for these simulated experimental sessions are as for Experiment 1a, with the following exceptions.

For these simulated experimental sessions the kurtosis of E.D.s was set to 3 (i.e. to an excess kurtosis of 0), and different levels of skew were sampled instead. E.D.s were either associated with 0 skew, or Category A inputs values (left tilted inputs) were positively skewed (by 0.16 or by 0.32), and Category B input values (right tilted inputs) were negatively skewed (by 0.16 or by 0.32). So, where simulated Experiments 1a and 1b sampled excess kurtosis levels of 0, 0.2 and 0.4, Experiments 2a and 2b will sample skew differences of 0, 0.32 and 0.64.

## Results

Confidence to perceptual slope ratios are plotted in Figure 4a, for experimental sessions that simulated a metacognitively ideal participant. As the simulated participant is metacognitively ideal (i.e. the same information has informed perceptual and confidence decisions and analyses), these ratios should cluster about a value of 1. As can be seen in Figure 4a, slope ratios do cluster about a value of 1, with nearly all falling within 5% of this value. These data therefore show that our new method of estimating metacognitive sensitivity is robust when analyses are informed by E.D.s that are differently skewed.

Confidence to perceptual slope ratios are plotted in Figure 4b – for experimental sessions where the simulated participant was metacognitively insensitive. When the simulated participant was metacognitively insensitive, slope ratios were all less than 1, with all but 1 falling more than 5% below this value.

## Simulated Experiment 2b: SDT-based analyses of confidence and skew

Details are as for Experiment 2b, with the following exceptions.

The kurtosis of both the Non-Target and Target E.D.s (see Figure 1) was set to 3 (i.e. to an excess kurtosis of 0, consistent with a normal distribution). Non-Target and Target E.D.s were either both associated with 0 skew, or Non-Target E.D.s (left tilted inputs) were positively skewed (by 0.16 or by 0.32), whereas Target E.D.s (right tilted inputs) were negatively skewed (by 0.16 or by 0.32).

## Results

Meta d’ to d’ ratios are plotted in Figure 4c, for experimental sessions that simulate a metacognitively ideal participant. As can be seen in Figure 4c, these ratios only broadly cluster about a ratio of 1 when the E.D.s informing data analyses were normally shaped (black data). When analyses were informed by E.D.s that have different skews, all ratios were > 5% lower than a ratio of 1. This shows that S.D.T. based analyses of confidence can encourage erroneous conclusions when analyses are informed by E.D.s that have different skews. Here the same information has informed d’ and meta d’ analyses, and yet results would encourage a researcher to conclude the participant had been metacognitively insensitive (i.e. that a different source of information has shaped confidence ratings).

Meta d’ to d’ ratios are plotted in Figure 4d, for experimental sessions that simulate a metacognitively insensitive participant. As can be seen in Figure 4d, these slope ratios are all substantially less than 1. This shows that SDT-based analyses of confidence are sensitive to cases of genuine metacognitive insensitivity, but these true detections of metacognitive insensitivity are qualitatively indistinguishable from the false detections of metacognitive insensitivity (i.e. meta d’ : d’ ratios < 1) depicted in Figure 4c. So, classifications of these participants as metacognitively insensitive are not diagnostic, as the same process has classified metacognitive ideal simulated participants as metacognitively insensitive (see Figure 4c).

## Discussion

Our two sets of simulated experimental sessions demonstrate that some popular SDT-based analyses of confidence will systematically underestimate metacognitive sensitivity when E.D.s are not normally shaped, either because they have excess kurtosis (see Figure 3), or because they are oppositely skewed (see Figure 4). Moreover, we have shown that our new, slope-based test of metacognitive sensitivity is more robust to violations of the normality assumption.

While it is interesting that our new slope-based test is conceptually more robust relative to a currently popular SDT based analysis of confidence (see Figures 3 & 4), we need to validate this approach for use in experiments with humans. We attempt this by re-examining an established relationship between the range of direction signals within a stimulus and perceptual confidence. When direction signals are generated by groups of moving dots, confidence is disproportionately undermined, relative to the precision of perceptual decisions, when there is a large as opposed to a small range of different direction signals to either side of the average direction (de Gardelle & Mammassian, 2015; Spence et al., 2016). The results of these studies were unclear as to whether the disproportionate impact of the range of direction signals was due to a decline in metacognitive sensitivity, or if this was due to a confidence bias. However, a subsequent study used a SDT-based analysis of confidence (Fleming, 2017), and found evidence that the effect is due to a conditional confidence bias (Spence et al., 2018). According to this evidence, participants would be equally able to distinguish between good and bad perceptual decisions across experimental conditions (i.e. they would have equal levels of metacognitive sensitivity), but they would be overall biased to report having lower confidence when judging the average direction of inputs that contain a larger range of direction signals.

Given our concerns regarding SDT-based analyses of confidence, we thought it was worth re-examining the relationship between decisional confidence and the range of direction signals (de Gardelle & Mammassian, 2015; Spence et al., 2016) in two behavioural experiments with human participants. These experiments will be matched in all details, other than on the reporting of confidence. In Experiment 3 people will use a continuous scale to report on confidence, whereas in Experiment 4 they will use a Likert scale to select one of a limited number of confidence ratings.

Another manipulation that can impact on measures of perceptual confidence is confirmation bias (e.g. Braun et al., 2018; Rollwage et al., 2020). When a preliminary perceptual decision is made, sampling of further perceptual evidence can become biased, with an amplification of decision congruent evidence, and insensitivity to decision incongruent evidence (Braun et al., 2018; Peters et al., 2017; Rollwage et al., 2020). To further validate our approach, in Experiment 5 we assess the impact of a confirmation bias, driven by having participants guess what stimulus will be presented in the subsequent trial.

To preface these results, in all our empirical experiments with human participants we replicate the findings of prior studies regarding metacognitive sensitivity, which were obtained using SDT-based analyses of confidence (Fleming and Lau, 2014; Maniscalco & Lau, 2012, 2016). In Experiments 3 and 4 we find that a wider range of direction signals impacts on confidence by encouraging a confidence bias, without any discernible impact on metacognitive sensitivity (as per Spence et al., 2018). In Experiment 5 we find that an encouraged confirmation bias reduces metacognitive sensitivity for bias-confirming evidence, while inducing a confidence bias for contradictory relative to confirming sensory evidence (as per Braun et al., 2018; Peters et al., 2017; Rollwage et al., 2020).

## Experiment 3: Direction signal range, Metacognitive Sensitivity and Confidence Bias

### Methods

A total of 26 volunteers participated, 16 female, with a mean age of 23 (S.D. 3.8). This would have delivered ∼0.8 power to detect the effect size reported in the original study with a specified alpha of 0.01 (Spence et al., 2015). Data recordings failed for 3 participants, so the final sample consisted of 23 volunteer participants, with a mean age of 23 (S.D. 3.4). This should still deliver ∼0.8 power to detect the effect size reported in the original study with a specified alpha of 0.05 (Spence et al., 2015). All participants were naïve as to the experimental hypotheses. Thirteen of the final group of volunteers participated in return for course credit. Ten of the final group of participants were compensated with $40 for their participation. All participants completed experimental sessions while seated in a dimly lit room, viewing stimuli from a distance of 57 cm with their head restrained by a chin-rest. The study was approved by The University of Queensland research ethics committee, and was conducted in accordance with the principles of the Declaration of Helsinki.

### Stimuli

Stimuli consisted of random dot kinematograms (RDKs), generated using a Cambridge Research Systems ViSaGe stimulus generator driven by custom Matlab R2013b (MathWorks, Natick, MA) software and presented on a gamma-corrected 19 inch Dell P1130 monitor (resolution: 1600 x 1200 pixels; refresh rate: 60 Hz). Each RDK consisted of 101 individual white dots, each subtending 0.2 degrees of visual angle (dva) in diameter at the retinae, and drifting in a linear direction at a speed of 5 dva / second. Dots were presented against a black background within a circular aperture with a diameter subtending 4 dva. The individual lifetime of each dot was 200ms, after which it was re-drawn at a random position within the aperture. Each dot was assigned a random initial age (between 0 and 200ms).

In the center of the test display, there was a small (diameter subtending 0.46 dva) red bull’s eye configuration that served as the fixation point, which participants were instructed to stare at throughout all stimulus presentations.

Individual dots within RDKs translated in a uniform range of different directions (+/- 5 ° or 20°) about an mean direction, which was slightly to the left or right of vertical. RDK presentations persisted for 1 second. Immediately after, participants were asked to simultaneously report on the average direction (left / right of vertical) and on their level of confidence (from a minimum rating of 5% on a linear scale, up to a maximum rating of 100%) by moving a mouse left or right, by an amount that was indicative of confidence. The mouse movements controlled the direction (left or right) a green bar was stretched into, and how far the green bar was stretched (which indicated a level of confidence along a continuous scale). Participants would press a mouse button to finish their combined report. On alternate trials, RDKs moved predominantly up or down, and slightly to the left or right of vertical by a magnitude that was controlled via our adaptive sampling routines.

### Matlab scripts used to setup, control and analyze experimental results

Our intention here is to provide a tutorial, describing how an experimenter can use the Matlab scripts we provide, as supplementary material and via our website <u>Metacognitive Sensitivity:</u> <u>Slope Test (uq.edu.au)</u>, to setup, control and analyze the results of an experiment that measures metacognitive sensitivity and confidence bias. Details are also pertinent for understanding this specific experiment.

### meta_SETUP.m

Adaptive sampling routines are set up by calling a custom Matlab m file, **meta_SETUP.m**. This file requires at least 4 variables to be specified prior to calling the script, with an option to specify 3 additional variables. The first variable, **N_Conditions** specifies the number of different experimental conditions. In our case, we set this to 2, as we wanted to have one condition where the range of direction signals was +/- 5° from an average direction, and a second condition where the range of direction signals was +/- 20°. The second essential variable is **Guess_Low_Asymptote**. This is the experimenters initial guess as to what test value will likely coincide with people making High-Confidence (confidence ratings that are greater than the participants’ overall average confidence rating across all trials) Category A responses (in this case, decisions indicating that the mean direction was to the left of vertical) on 90% of trials. We set **Guess_Low_Asymptote** to -10°. It is not essential that this estimate be accurate, as sampling is dynamic and will adjust. The third essential variable is **Guess_High_Asymptote**. This is the conceptual mirror opposite of **Guess_Low_Asymptote**, the experimenters first guess of the test value that will almost always result in High-Confidence Category B responses. The fourth essential variable is **N_Trials_Per_Condition**. This specifies how many trials the experimenter wants to run per experimental condition. We set this to 200, and would not recommend a lesser number than this. The final essential variable is **Max_Possible_Confidence_Rating**. This specifies the maximum possible confidence rating a participant can give. In this experiment, people rated confidence along a continuous scale, from which we calculated a proportional confidence rating. So, we set this variable to 1.

With these five essential variables specified, a call to **meta_SETUP.m** will **1)** generate a variable **Min_Step_Size**, that specifies by what magnitude test values will be adjusted according to participant responses. **Min_Step_Size** is also one of 3 optional variables that an experimenter can set if, for instance, they need to ensure test values are adjusted in a step size that can be sampled by the testing apparatus. If this is not specified before calling **meta_SETUP.m**, it will be set to 20% of the range separating **Guess_Low_Asymptote** and **Guess_High_Asymptote**. In our case we did not specify this variable, so **Min_Step_Size** was set to 4° by the experimental code.

A call to **meta_SETUP.m** will also, **2)** generate a variable **Staircase_Sample_Pts** which sets initial test values that will be sampled by each of 5 staircase procedures that will be interleaved for each test condition. The first and fifth of these are set to the first guess asymptotes, the third is set to the mid-point between first guess asymptotes, and the second and fourth are set to mid-points in-between each first guess asymptote and the overall initial midpoint. A call to **meta_SETUP.m** will also, **3)** generate a variable **N_Trials**, and set this to **N_Conditions * N_Trials_Per_Condition**. Experimenters should use **N_Trials** to control the experimental loop that controls trial data collection (but they should not set this parameter in their own scripts, to avoid inadvertent conflicts). A call to **meta_SETUP.m** will **4)** generate a variable **Condition_N** and set this to a number that specifies what experimental condition the experimenter should sample on the first trial of the experiment. Finally, a call to **meta_SETUP.m** will **5)** generate a variable **Test_Val** that specifies what test value should be sampled by the experimenter on the first trial of the experiment. As readers can see in provided code, **meta_SETUP.m** conducts many more operations, but we have specified the 5 major operations that are needed to set up an experiment.

### meta_UPDATE.m

On each trial, the experimenter should record **1)** a perceptual response, stored as a variable **Perceptual_Resp**. This should either be set to a value of 1 (to indicate a Category A response, in this case a response indicating that the participant thought the average direction was to left of vertical) or 2 (a Category B response). The experimenter should also record **2)** a confidence rating, saved as a variable **Confidence_Rating**. To accommodate Likert scales, this can take on any value. In this experiment, we set **Confidence_Rating** to a value between 0 and 1 as participants made confidence ratings by selecting a point along a continuous scale, which were expressed as a proportional confidence rating. Having specified **Perceptual_Resp** and **Confidence_Rating** the experimenter should then call **meta_UPDATE.m**. This script will update trial records, and assign new values to **Condition_N** and to **Test_Val**, to specify what experimental condition and test value should be sampled on the next trial. The script **meta_UPDATE.m** requires an estimate of where a participants’ confidence boundaries might be, to appropriately target sampling. To achieve this, during experimental sessions confidence ratings on each trial are categorized as low or as high relative to an evolving partition value, *v*:

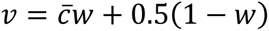

where 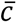 is the mean confidence expressed up to *t*, the current trial. Mean confidence is given linearly increasing weight, *w*, over the first 21 trials, according to:

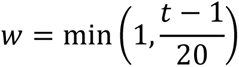

Hence confidence ratings are initially categorized using a partition value set to half the maximal possible confidence rating, but this value soon converges to the mean confidence rating expressed so far. Note this categorization is performed purely to facilitate targeted samplings of test inputs. It does not play a role in subsequent data analyses.

### meta_ANALYSE.m

When the experiment finishes, the experimenter should call **meta_ANALYSE.m**. This script requires that an array called **Exp_Results** has been created. This is an array with a row for each experimental trial and 4 columns, with trial condition numbers stored in the first column, test input values in the second, perceptual category responses in the third and confidence ratings in the fourth. This array is created and updated by **meta_UPDATE.m**, but experimenters could setup their own version of this array and use **meta_ANALYSE.m** to assess their data should they wish to bypass **meta_UPDATE.m**.

When executed, **meta_ANALYSE.m** will sort trial data into experimental conditions, and fit cumulative Gaussian functions to **1)** The proportion of Low-Confidence Category A + any Category B responses, as a function of tested values. We refer to this as the *Category A Confidence function* (see blue data, Figure 2), **2)** The proportion of Category B responses, as a function of tested values. We refer to this as the *Perception function* (see black data, Figure 2), and **3)** The proportion of High-Confidence Category B responses, as a function of tested values. We refer to this as the *Category B Confidence function* (see red data, Figure 2). In each case, whenever the code fits a cumulative Gaussian function to data, it also calculates residuals for datapoints relative to the average Y-axis value of each dataset (to represent chance responding). If the set of chance responding residuals is not 1.5 times greater than the set of residuals that relate to the function fit, no function fit statistics are recorded, and the experimenter is warned.

From function fits, the script generates **1)** a variable **Confidence_Crit** that is set to the average X-axis offset of the confidence functions from the Perception function, **2)** A variable **mSlopeRatios** that is set to the average slope of fitted confidence functions at their inflection points to the slope of the Perception function at its inflection point. For each of these variables a value is generated for each experimental condition.

Of the output variables, **Confidence_Crit** indicates what magnitude of offset, from the perceptual category boundary, a participant requires before they will endorse a perceptual category decision with a relatively high-level of confidence, and **mSlopeRatios** estimates how precise confidence ratings are relative to perceptual decisions. If a participant is metacognitively ideal, **mSlopeRatios** should be equal to 1. If confidence ratings are shaped by additional information **mSlopeRatios** should be equal to some value less than 1. The script also outputs **meta_Difference**, which gives the difference between the average slope of fitted Confidence functions at their inflection points and the slope of the Perception function at its inflection point. If a participant is metacognitively ideal, **meta_Difference** should be equal to 0. If confidence ratings are shaped by additional information, **meta_Difference** should be equal to same value less than 0.

## Results

The results of Experiment 3 are depicted in Figure 4. Metacognitive slope ratios (Avg. Confidence Slope to Perceptual Slope) were less than 1 for both of our experimental conditions (see Figure 4a), which indicates that confidence ratings were shaped by different information, relative to perceptual category decisions. There was, however, no evidence for a robust difference between our two conditions, with a Bayes factor analysis revealing moderate evidence for the null hypothesis, that there would be no difference in Avg. Confidence to Perceptual slope ratios across our two experimental conditions (see Figure 4a).

Metacognitive slope relationships are also expressed as conditional differences (Avg. Confidence Slope - Perceptual Slope, see Figure 4b). Average differences for both of our experimental conditions were < 0, suggesting that confidence ratings were shaped by different information relative to perceptual category decisions. There was, however, no evidence for a robust difference between levels of metacognitive sensitivity across two conditions, with a Bayes factor analysis revealing moderate evidence for the null hypothesis, that there would be no conditional difference (see Figure 4b).

The results of Confidence criterion analyses are depicted in Figure 4c. People adopted a more conservative confidence criterion (they needed a greater average direction offset from vertical to report a relatively high level of confidence) for RDKs with a broader range of directions signals (see Figure 4c). A Bayes factor analysis revealed moderate level of evidence for the alternative hypothesis, that people would adopt different confidence criteria when judging the average direction of RDKs that had different ranges of direction signals. Overall, these data suggest that the impact of a larger range of direction signals on confidence is to cause people to be more conservative when deciding if they should endorse their perceptual decisions with a relatively high-level of confidence. These data suggest that there is no adverse impact on metacognitive sensitivity, on the ability of the participant to distinguish between relatively good and bad direction decisions.

In Experiment 3, people made simultaneous perceptual direction decisions and confidence ratings, by choosing a direction in which to stretch a bar (left or right) to indicate the average test direction, and by choosing how far to stretch that bar (to indicate a level of confidence along a continuous scale) before they pressed a button to finish their setting. Many studies use a sequential report instead, with people making a perceptual decision before they give a decisional confidence rating. Many studies also use a Likert scale for reporting on confidence. We wanted both to see if these changes might undermine metacognitive sensitivity when tests have a larger range of direction signals, and we wanted to validate our approach to setting up, controlling, and analyzing the results of a more typically experimental design.

## Experiment 4: Direction signal range, Metacognitive Sensitivity and Confidence Bias measured using a Likert scale and sequential decisions

Almost all details of Experiment 4 were as for Experiment 3. The same volunteers participated using the same apparatus, and the experiment was controlled using the same Matlab scripts as we have described for that experiment. The only substantive difference was that immediately after test presentations, people were asked to press one of two buttons to indicate if the mean test direction had been to the left or right of vertical. After this response, they were then asked to rate their level of confidence in their direction decision, by using a mouse to highlight and click on one of 5 confidence ratings labelled ‘1 (Guessing’, ‘2’, ‘3’, ‘4’, or ‘5 (Certain)’.

## Results

The results of Experiment 4 are depicted in Figure 5. Metacognitive slope ratios were less than 1 for both experimental conditions (see Figure 5a), suggesting that confidence ratings were shaped by different information, relative to perceptual category decisions. Again, there was no evidence for a robust difference between conditions in terms of metacognitive sensitivity, with a Bayes factor analysis revealing moderate evidence for the null hypothesis, that there would be no conditional difference in metacognitive slope ratios (see Figure 5a).

**Figure 5.**
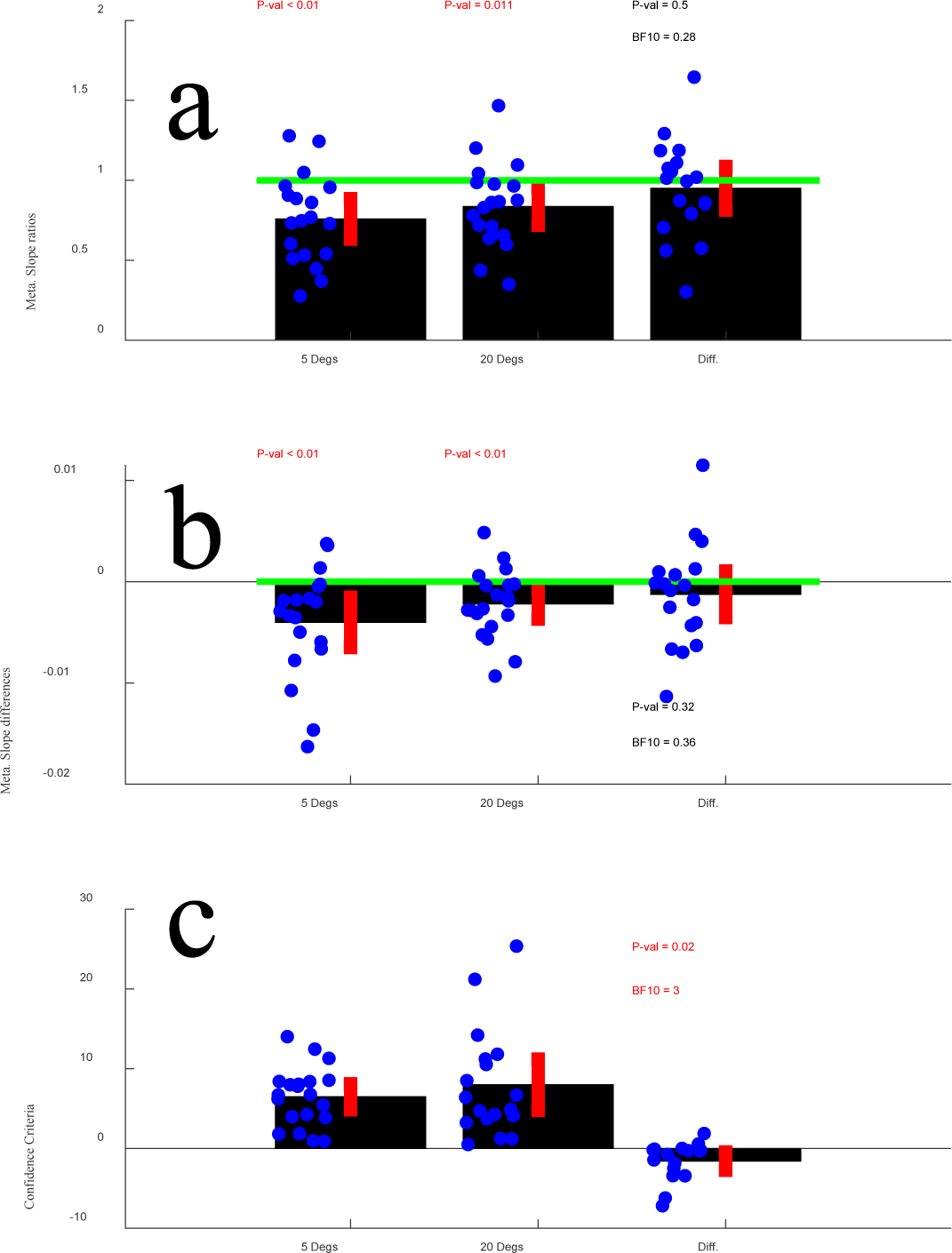
Results of Experiment 4. All details regarding this figure are as for Figure 4.

As per Experiment 3, metacognitive slope relationships are also expressed as conditional differences (see Figure 5b). Both average conditional differences were < 0, suggesting that confidence ratings were shaped by different information relative to perceptual direction decisions. There was, however, no robust evidence for there being a difference between these conditional differences (see Figure 5b).

The results of Confidence criterion analyses are depicted in Figure 5c. Again, people adopted a more conservative confidence criterion for tests with a broader range of directions signals (see Figure 5c), with a Bayes factor analysis revealing moderate level of evidence for the alternative hypothesis, that people would adopt different confidence criteria when judging the average direction of tests that had different ranges of direction signals.

The results of Experiment 4 reiterate those of Experiment 3, reinforcing the view that the impact of a larger range of direction signals on confidence is to cause people to be more conservative when deciding if they should endorse their perceptual category decisions with a relatively high-level of confidence.

The results of Experiments 3 and 4 both conceptually replicate the results of an earlier study, which had suggested that a larger range of direction signals would encourage people to be more conservative when deciding if they should endorse their perceptual direction decisions with a relatively high level of confidence (Spence et al., 2018). In that study, and in our Experiments 3 and 4, there was no robust evidence for a detrimental impact on metacognitive sensitivity from having a larger range of direction signals in tests. We also wanted to validate our approach to setting up, controlling, and analyzing the results of a confidence experiment in a context where other studies had suggested a conditional difference in metacognitive sensitivity. Experiment 5 was therefore designed to encourage a confirmation bias, by having people guess at what direction of input they would see in a subsequent test.

## Experiment 5: Direction Perception, Confidence and Confirmation Bias

Almost all details for Experiment 5 are as for Experiment 3. The same volunteers participated using the same apparatus, and the experiment was controlled using the same Matlab scripts as we described for Experiment 3. The only substantive differences were that all tests contained a range of direction signals +/- 30° from the test mean, and prior to test presentations people were asked to guess what they thought the next average test direction would be. They were encouraged to consider their last 3 guesses (Left or Right), in order to help them make a prediction. A record of these was displayed, initialized with place-holding ‘none’ labels which were progressively filled as trials were completed and predictions made. Our hope was to elicit a gambler’s fallacy-like (e.g. Xu & Harvey, 2014) sense that a participant could anticipate these randomized stimulus presentations by considering the pattern of recent presentations. Experimental conditions refer to tests that were Consistent with guesses, or Inconsistent with guesses.

## Results

The results of Experiment 5 are depicted in Figure 6. In this case metacognitive slope ratios were not, on average, less than 1 (see Figure 6a). Nor was there any robust evidence for a difference between conditions in terms of metacognitive sensitivity (see Figure 6a). Metacognitive slope relationships are also expressed as conditional differences (see Figure 6b). In this analysis there is evidence that Confidence function slopes were shallower than Perceptual function slopes when tests were Consistent with guesses, whereas there was no evidence of this difference for tests that were Inconsistent with guesses. There was also some evidence for a conditional difference in these estimates of metacognitive sensitivity, with evidence of a relatively reduced sensitivity when tests were consistent with guesses.

**Figure 6.**
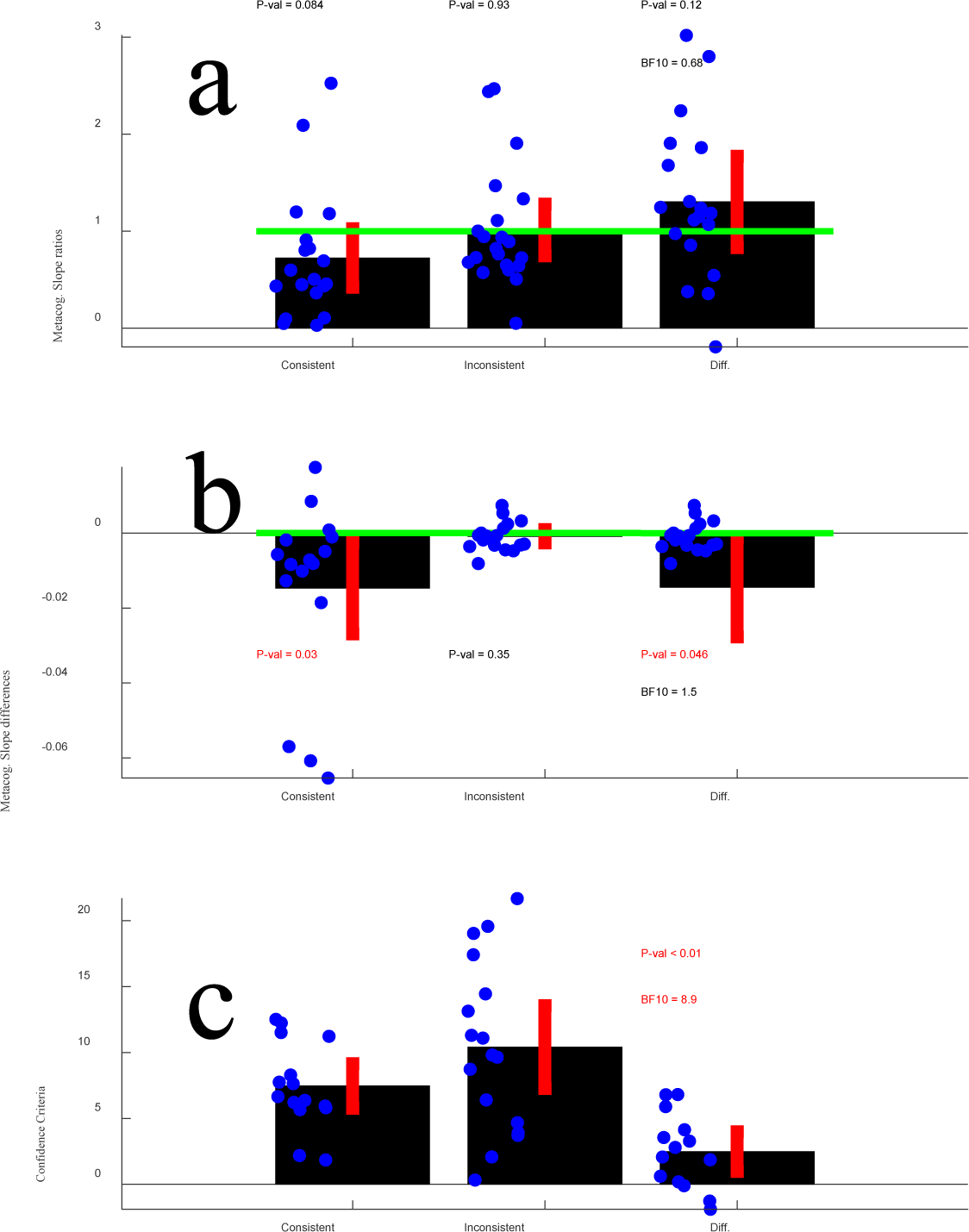
Results of Experiment 5. All details regarding this figure are as for Figure 4, except that experimental conditions refer to tests that were Consistent with peoples guesses, or were Inconsistent.

The results of Confidence criterion analyses are depicted in Figure 6c. These data show that people adopted a more conservative confidence criterion for tests that were Inconsistent with people’s guesses as to what test direction would be presented (see Figure 6c). A Bayes factor analysis revealed a moderate level of evidence for the alternative hypothesis, that people would adopt different confidence criteria when judging the average direction of tests that were Consistent or Inconsistent with their guesses as to what test direction would be presented.

## General Discussion

We have described a new method to estimate metacognitive sensitivity and confidence criteria. In simulations, we have shown this method (see Figures 2-3) is more robust to violations of the normality assumption than an established SDT-based analysis (Fleming and Lau, 2014; Maniscalco & Lau, 2012, 2016). We then validated the new method in a series of behavioural experiments with human observers. Our data from these experiments suggest that having a broader range of direction signals causes people to be more conservative when rating their confidence in decisions about the average direction (a confidence bias effect, see Figures 4-5). Metacognitive sensitivity (their ability to distinguish between likely right and likely wrong decisions) is undiminished (see Figures 4-5, also see Spence et al., 2018). In an experiment looking at the impact of conformation bias, we find evidence this can both selectively reduce metacognitive sensitivity for decisions regarding expected inputs (see Figure 6), and cause people to be more conservative when rating their confidence in decisions regarding expected inputs, as opposed to unexpected inputs (see Figure 6).

### What is wrong with some popular SDT-based analyses of confidence?

Currently popular SDT-based analyses of confidence (Fleming and Lau, 2014; Maniscalco & Lau, 2012, 2016) make a backwards inference from proportions of correct and incorrect high-confidence perceptual decisions. The inference is an estimate of how separated distributions of the different experiences triggered by repeated presentations of each of two inputs must be, to explain proportions of correct and incorrect high-confidence decisions. To achieve this, an assumption is made regarding what shape E.D.s have. While SDT as a framework is agnostic about the precise shape of these distributions (Green & Sweets, 1966), implementations of the theory often assume a normal shape. Any difference between the assumed and actual shape of the two E.D.s can result in marked differences between expected and actual proportions of high-confidence decisions.

To clarify, imagine repeated presentations of a physically vertical input. Current implementations of SDT used to measure confidence (Fleming and Lau, 2014; Maniscalco & Lau, 2012, 2016) assume a person will have different experiences of that input, most often seeing it as vertical, but also often as slightly left and right tilted, and very occasionally as considerably left and right tilted. The overall shape of the distribution describing these different experiences is assumed to be normal (Gaussian). If, instead, the actual shape was non-normal, because it had excess kurtosis (fat tails), then there would be more experiences of considerable left and right tilt than would be predicted by a normal distribution. A backwards inference informed by these greater numbers of extreme experiences would result in a systematic error, in an inference that the two assumed distributions are closer together than they are in fact, to account for the greater numbers of high-confidence decisions than are predicted by a normal distribution. This is a substantive issue for popular SDT-based analyses of confidence (Fleming and Lau, 2014; Maniscalco & Lau, 2012, 2016), as this situation is interpreted as evidence of metacognitive insensitivity – as evidence that confidence has been shaped by a different information relative to perceptual category decisions. However, this outcome can also follow from E.D.s having had a non-normal shape (see Arnold et al., 2023; Miyoshi et al., 2022).

This impact of any distortion in the shape of E.D.s from an assumed normal will tend to scale with how conservative a person is when deciding if they should report having confidence in a category decision. To illustrate, in Figure 7a we have plotted a normal distribution (black) and a fat-tailed distribution with the same specified mean and s.d. but excess kurtosis (0.4, red). In Figure 7b, we plot the proportion of the fat-tailed distribution that lies to the right of each X-axis value / the proportion of the normal distribution that lies to the right of each X-axis value. Note that as you near the right-side tail of these distributions, there is a far greater proportion of the fat-tailed distribution remaining than would be expected if the distribution had a normal shape. Bear in mind that these distributions represent the numbers of trials that result in different experiences. A backwards inference assuming normality based on the proportion of responses represented by the tails of the fat-tailed distribution will therefore mistakenly assume that this distribution is fatter than it is, and is therefore less separated from a mirror opposite distribution in S.D. units. This is why popular SDT-based analyses of confidence (Fleming and Lau, 2014; Maniscalco & Lau, 2012, 2016) fail when confronted with excess kurtosis, particularly when a simulated participant is conservative when deciding if a perceptual decision should be endorsed with relatively high confidence (see Figure 3).

**Figure 7.**
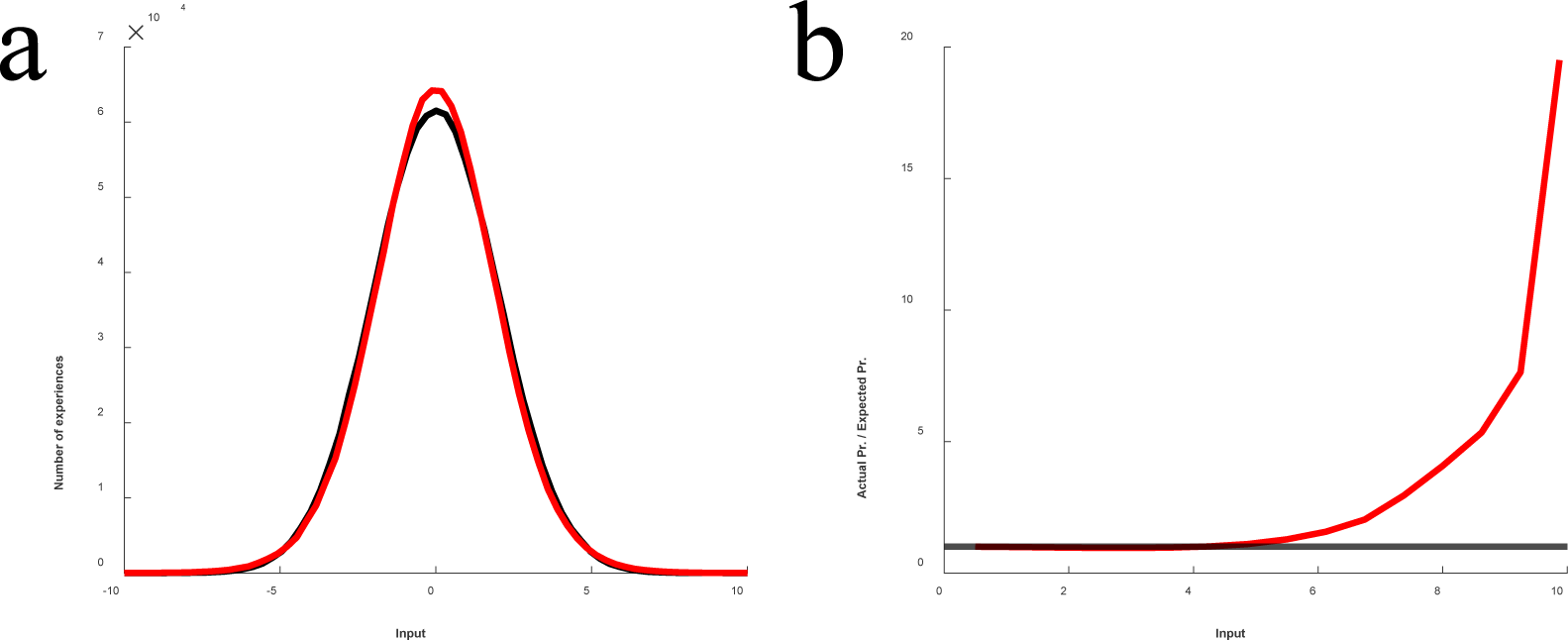
**a)** Plot of a normal distribution (black) and a fat-tailed distribution (red) with the same specified mean and standard deviation, but with an excess kurtosis of 0.4. **b)** Plot of the proportion of the fat-tailed distribution to the right of each X-axis value / the proportion of the normal distribution to the right of each X-axis value. This illustrates that as you near the right-side tail of the fat distribution, there is a greater proportion of the fat-tailed distribution remaining, relative to what would be expected of a normal distribution. In a SDT-based experiment, this would be analogous to there being a greater proportion of high-confidence decisions than you would expect if the underlying distribution had a normal shape.

We have chosen to illustrate dilemmas for SDT-based analyses of confidence with a detailed description of how these relate to a distribution with excess kurtosis. The issues for skewed distributions are, however, very similar. In essence, greater or lesser proportions of trials are located at the tails of skewed distributions than is predicted by the normality assumption, and this can cause SDT-based analyses of confidence that assume normality to over or underestimate meta d’ relative to d’, even when these statistics are calculated from a common dataset (see Figure 4, and Arnold et al., 2023).

### Why is our method more robust?

Our method involves fitting cumulative Gaussian functions to behavioural data, and conceptually this analysis commits to the normality assumption in precisely the same way as many other SDT-based analysis (for a related discussion, see Yarrow et al., 2011). So why is our approach more robust?

In our approach, we fit cumulative Gaussian functions to data sampled from many more than just 2 test inputs (see Figure 2). These function fits assume that each datapoint is derived from its own Gaussian shaped ED (for a related discussion, see Yarrow et al., 2011). For the individual datapoints, the deviance between outcomes suggested by analyses informed by normally and by non-normally shaped EDs will depend on the positioning of the ED relative to the relevant criterion (be that a perceptual decision, or a confidence criterion). The closer the ED is to the relevant criterion, the less the difference in outcomes. This relationship is evident in Figures 3 – 4 and in Figure 7. As depicted in Figure 7b, the reason is that there is little difference in the proportion of an ED positioned to either side of a criterion value, regardless of the precise shape of the ED, when the ED is nearly centred on a criterion value. So, a key reason our approach is more robust against violations of the normality assumption, relative to currently popular SDT-based analyses (Fleming and Lau, 2014; Maniscalco & Lau, 2012, 2016), is that by sampling more than 2 inputs we ensure function fits are constrained by some datapoints that are derived from EDs that are proximate to a relevant criterion value.

Our slope test for metacognitive sensitivity has some features in common with another recent approach to measuring metacognitive sensitivity. Mammassian & Gardelle (2022) have implemented an approach, grounded in SDT (as is ours) but augmented by use of ideal observer analyses and modelling (Barlow, 1962; Geisler, 1989). In common with our approach, this method involves sampling a reasonable number of different inputs and fitting psychometric functions. We therefore suspect it will prove similarly robust to violations of the normality assumption. At this point, Mammassian & Gardelle’s (2022) ideal observer modelling approach is specifically targeted at studies where participants must choose which of their last 2 perceptual decisions evoked greater feelings of confidence. In addition, they implement a detailed generative model that describes *how* feelings of confidence might arise and, given a sufficiently sized dataset, allows for factors that shape confidence to be considerably fractionated. By contrast, our approach is targeted at common study designs, and is primarily concerned with estimating two key statistics – metacognitive sensitivity and confidence criterion. We would hope the two approaches will prove complementary. Our approach might benefit researchers who want to implement a robust measure of metacognitive sensitivity in a simple study design. Mammassian & Gardelle’s (2022) approach should benefit researchers who wish to conduct a more demanding modelling-based study and analyses of data, in order to partition of the influences of confidence in detail.

### Things to be aware of if you use our method

Our method is easy to implement in standard experimental designs. We have also provided code designed to make this implementation easy. We hope this will be useful to other researchers. There are, however, issues you should be aware of.

Our approach requires a sampling of inputs that result in people reliably making High-Confidence Category A responses at one extreme of the tested range, and High-Confidence Category B responses at the other extreme, with intervening test values resulting in many variable Low-Confidence category decisions. If there is no variance in a participants’ confidence ratings, our approach will fail.

We would suggest it is worth conducting a little training to screen participants prior to formal data collection (this process and exclusion criteria should be pre-registered). A participant could be shown exemplar inputs that should be trivial to categorise (say stimuli that are tilted +/- 30 degrees from vertical, or some equivalent for your experimental design). The experimenter would need to confirm that each participant can reliably categorise these exemplars, and if they can, they could then instruct the participant that if these inputs can clearly be discerned, they should report a relatively high-level of confidence. Participants could also be informed that other inputs will be ambiguous, and that these should evoke a confidence rating that corresponds with the ambiguity of the input.

Without such training, we have found many participants will never reliably endorse perceptual category decisions with high-confidence. This might be because they have failed to understand task instructions, or because they think there is some unknown trick to the experiment that undermines their confidence. Regardless of the reason, if people will not reliably endorse a blatantly obvious perceptual category decision with a relatively high level of confidence, our approach will fail. But perhaps in a good way. Our data plots will reveal when a participant has failed to understand task instructions, whereas this can be less obvious using other approaches.

### What do our data say about confidence in direction decisions?

Our experiments were primarily conducted to validate our new method of estimating metacognitive sensitivity and confidence bias, through behavioural experiments with human participants. Our data do, however, allow us to make some observations.

First, we have found evidence that a wider range of direction signals impacts on confidence by encouraging participants to be more conservative when deciding if they should endorse a perceptual category decision with relatively high confidence (see Figures 4-5). We found no evidence for an impact on metacognitive sensitivity. These findings replicate those of an earlier study that used a different measure of the impact of confidence (Fleming, 2017) to reach the same conclusions (Spence et al., 2018). We have also found evidence that an encouraged confirmation bias can reduce metacognitive sensitivity for bias-confirmatory evidence, and induce a confidence bias for confirmatory relative to contradictory sensory evidence (as per Braun et al., 2018; Peters et al., 2017; Rollwage et al., 2020). In each case, our data have provided convergent evidence in relation to existing findings.

### Conclusions

We have described a novel means of measuring metacognitive sensitivity and confidence crtiteria. This method is more robust against violations of the normality assumption than currently popular SDT-based analyses of confidence (e.g. Fleming, 2017; Fleming and Lau, 2014; Maniscalco & Lau, 2012, 2016). We provide code you can access to implement our procedures (https://www.psy.uq.edu.au/∼uqdarnol/MetaCog_Slope_Test.html).

## Acknowledgements

This research was supported by a Discovery Project Grant DP200102227, funded by the Australian Research Council, awarded to D.H.A.

## Data and materials availability

All data and analysis scripts for this project will be made available via UQeSpace https://espace.library.uq.edu.au

## Supplemental note on the formulation of the slope test for metacognitive sensitivity

### Notations

Let *Y* = *F*(*X*; γ, λ, μ;, σ) be a psychometric function that describes the probability that an observer will give a category response of 1 (in the context of a forced choice categorization task) to a stimulus input with value *X*. Specifically, we define the function as follows

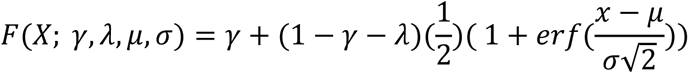

where γ is the probability that the observer will give a category response of 1 when *X* → −∞ (i.e., the “guess rate”), and λ is the probability that the observer will give a category response of 0 when *X* → +∞ (i.e., the “lapse rate”), μ; is the stimulus value at which the p(response = 1) = 0.5 (i.e., the mean of the underlying cumulative normal distribution), σ is the standard deviation of the underlying normal distribution, and *erf*(*x*) is the error function

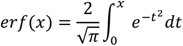

Then, *y* = *F*′(*X*; γ, λ, μ;, σ) denotes the slope of the function with respect to X (i.e., the probability density function of the normal distribution), and *F*′(*x*_0_; γ, λ, μ;, σ) denotes the slope of the cumulative normal distribution function at *x* = *x*_0_ given mean μ; and SD σ.

Under the SDT framework, the internal representation of the stimulus is corrupted by noise. For a stimulus with value *x* (e.g., stimulus orientation, motion direction, etc.), let 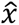 be the value of the noisy representation of the stimulus.

Here, we focus on the **decision process** of the SDT, which concerns the criteria.

Suppose an observer’s perceptual task is to categorize the stimulus as a target *T* or non-target *N*. SDT assumes there is a **perceptual criterion** *c*_0_ which the observer compares the noisy representation 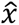 to commit to a perceptual category decision.

In addition to the perceptual task, suppose the observer has to rate their confidence in the quality of their perceptual decision. For simplicity, we assume this confidence response is binary (i.e., either low or high).

For each of the *T* and *N* stimulus categories, the SDT assumes there is a **confidence criterion** that separates high from low confidence responses. Let *c*_*T*_ be the confidence criterion for the *T* category and *c*_*N*_ be the confidence criterion for the non-target category. Assuming targets have a larger internal representation value than non-targets, the three criteria should go from smallest to largest as *c*_*N*_, *c*_0_, and *c*_*T*_.

Suppose the observer views a set of *n* stimuli *X* = {*x*_1_, *x*_2_, . . ., *x*_*n*_}, each of which is chosen from a set of *m* levels *L* = {*l*_1_, *l*_2_, . . ., *l*_*m*_}, with *m* being the number of fixed levels and *n* being the number of trials in the experiment. The *L* values correspond to stimulus values such as the orientation of gratings, the average motion direction of an RDK, etc..

Let *k*_*j*_ be the number of trials on which stimulus at level *l*_*j*_ is presented (e.g., *k*_1_ stimuli are presented with stimulus value *l*_1_) during the experiment. To each stimulus, observers give one response *Y* = {*y*_1_, *y*_2_, . . ., *y_m_*}, where each of the y is one of the following four response categories: {“High-confidence, Non-target”, “Low-confidence, Non-target”, “Low-confidence, Target”, “High-confidence, Target”}.

### Formulation of the slope test

In summary, responses are recoded with respect to decision criteria, and then fitted with a psychometric function (a cumulative normal distribution function here). This produces a confidence slope for two confidence criteria, and a perceptual slope for the perceptual criterion, the ratio between which is computed as a measure of metacognitive sensitivity.

| Range of $\hat{x}$ | Response | Recoding responses with respect to... | | |
| --- | --- | --- | --- | --- |
| | | Non-target confidence criterion $c_N$ | Perceptual criterion $c_0$ | Non-target confidence criterion $c_T$ |
| $\hat{x} < c_N$ | High-confidence, Non-target | 0 | 0 | 0 |
| $c_N \leq \hat{x} < c_0$ | Low-confidence, Non-target | 1 | 0 | 0 |
| $c_0 \leq \hat{x} < c_T$ | Low-confidence, Target | 1 | 1 | 0 |
| $\hat{x} \geq c_T$ | High-confidence, Target | 1 | 1 | 1 |

The procedure is described in the following parts.

### Part 1: Estimating the slope based on the confidence criterion for the non-targets

A. Recode the responses *Y* and define *Y*_*N*_ = {*y*_*N*1_, *y*_*N*2_, . . ., *y*_*Nn*_} as follows:

*y*_*N*i_ = 0 if *y*_i_ *X* “High-confidence, Non-target“;

*y*_*N*i_ *X* 1 otherwise.

For each fixed level of stimulus value *l*_*j*_, we bin the *k*_*j*_ responses to compute *p*_*Nj*_, which is the proportion of responses in which *y*_*N*i_ = 1.

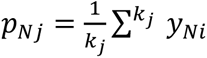 for all *y_Ni_* with the corresponding *x_i_* = l*_i_*

We then obtain a set of response proportions *P*_*N*_ = {*p*_*N*1_, *p*_*N*2_, . . ., *p*_*Nm*_} that reflect response categorizations for each stimulus level in *L* with respect to the criterion *c*_*N*_.

We then fit the cumulative normal distribution function to the *mm* data points (*L*, *P*_*N*_), with a set of weights *W* = {*w*_1_, *w*_2_, . . ., *w*_*m*_} given to each data point based on the number of trials for each level

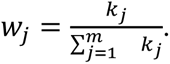

Formally, we estimate 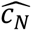 *and* 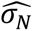 that best fit the function (with weights *W)*

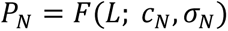

Afterwards, we compute the slope at the inflexion point (i.e., at the mean of the cumulative normal distribution function) of the fitted function by computing

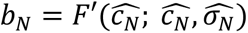

*b*_*N*_ represents the estimated sensitivity of the *c*_*N*_ criterion in separating “high-confidence, Non-target” responses with other responses.

#### Part 2: Estimating the slope based on the confidence criterion for the targets

A. We repeat every step in Part 1, but at 2A, we define *Y*_*T*_ = {*y*_*T*1_, *y*_*T*2_, . . ., *y*_*Tnn*_} as follows:

*y*_*TN*_ = 1 if *y*_*N*_ = “High-confidence, Target“;

*y*_*TN*_ = 0 otherwise.

B. Next, we go through the same steps in Part 1 to obtain the proportion of responses *P*_*T*_ based on *Y*_*T*_ and then perform the fitting to obtain 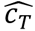 and 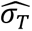

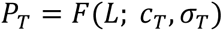

C. Then, we compute the slope at the inflexion point for this fit:

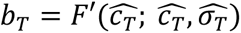

*b*_*T*_ represents the estimated sensitivity of the *c*_*T*_ criterion in separating “high-confidence, Target” responses with other responses.

#### Part 3: Estimating the slope based on the perceptual criterion

A. We repeat every step in Part 1, but at 3A, we define *Y*_0_ = {*y*_01_, *y*_02_, . . ., *y*_0*n*_} as follows:

*yy*_0i_ = 1 if *y*_i_ = “High-confidence, Target“ or “Low-confidence, Target”;

*y*_0i_ = 0 otherwise.

B. Next, we go through the same steps in Part 1 to obtain the proportion of responses *P*_0_based on *Y*_0_ and then perform the fitting to obtain 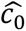 and 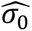

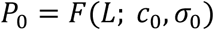

C. Then, we compute the slope at the inflexion point for this fit:

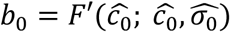

*b*_0_ represents the estimated sensitivity of the *cc*_0_ criterion in separating “Non-target” and “Target” responses.

#### Part 4: Computing the slope ratio

A. We first compute the confidence slope *bb*_*ccccnncc*_ as the average of the two slopes obtained in Parts 1 and 2

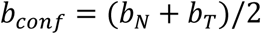

A. Then, the slope ratio *r* is defined to be the ratio between the confidence-criterion slope and the perceptual-criterion slope,

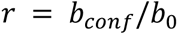

